# Cooperative Motility, Force Generation and Mechanosensing in a Foraging Non-Photosynthetic Diatom

**DOI:** 10.1101/2023.03.27.533254

**Authors:** Peng Zheng, Kayo Kumadaki, Christopher Quek, Zeng Hao Lim, Yonatan Ashenafi, Zhi Ting Yip, Jay Newby, Andrew J. Alverson, Yan Jie, Gregory Jedd

## Abstract

Diatoms are ancestrally photosynthetic microalgae. However, some underwent a major evolutionary transition, losing photosynthesis to become obligate heterotrophs. The molecular and physiological basis for this transition is unclear. Here, we isolate and characterize new strains of non-photosynthetic diatoms from the coastal waters of Singapore. These diatoms occupy diverse ecological niches and display glucose-mediated catabolite repression, a classical feature of bacterial and fungal heterotrophs. Live-cell imaging reveals deposition of secreted extracellular polymeric substance (EPS). Diatoms moving on pre-existing EPS trails (runners) move faster than those laying new trails (blazers). This leads to cell-to-cell coupling where runners can push blazers to make them move faster. Calibrated micropipettes measure substantial single cell pushing forces, which are consistent with high-order myosin motor cooperativity. Collisions that impede forward motion induce reversal, revealing navigation-related force sensing. Together, these data identify aspects of metabolism and motility that are likely to promote and underpin diatom heterotrophy.

## Introduction

Eukaryotes fall into fundamentally distinct groups based on their means of energy acquisition. Photoautotrophs— land plants and algae—derive energy from sunlight. By contrast, heterotrophs, such as animals and fungi, obtain energy by feeding on primary producers or each other. Mixotrophs combine these two strategies.

Eukaryotic photoautotrophs evolved several times through endosymbiosis between a heterotrophic eukaryote and photosynthetic microbe. Such an association between the ancestor of the Archaeplastida and a cyanobacterium led to the emergence of land plants, green and red algae, and glaucophytes. Afterwards, distinct algal lineages arose through secondary and tertiary endosymbiotic events, where the photosynthetic capacity was acquired from a eukaryotic red or green alga^1–4^.

The stramenopiles (also known as heterokonts) obtained their plastid through a secondary or higher endosymbiotic event with a red alga^5^. This group includes oomycetes, multicellular brown algae and unicellular diatoms^6^. With an estimated 100,000 species^7^, diatoms are one of the most abundant and diverse group of marine and freshwater microalgae^8–12^. Notably, they employ mineralization to construct silica-based^13,14^ cell walls (frustules) that fit together like the two halves of a petri dish. Diatoms possess either radial (centric diatoms) or bilateral (pennate diatoms) symmetry. A group of pennate diatoms - evolved a fine longitudinal slit through the frustule known as the raphe. These raphid pennate diatoms can move using a lineage-specific form of gliding motility^15,16^, and have undergone substantial evolutionary radiation to comprise the most species-rich and diverse lineage of diatoms^9^.

The raphe acts as a channel for secretion of a complex mixture of proteins and glycoproteins^17–20^ known collectively as extracellular polymeric substances (EPS). Motility can be blocked by an antibody to EPS^17^ and actin inhibitors^21^, suggesting that both play essential roles. Actin filaments occur in two prominent bundles that underlie the raphe just adjacent to the plasma membrane^22^. This arrangement supports a model of motility where myosin motors exert pushing forces on the extracellular EPS through a transmembrane protein^23^. However, this protein has yet to be identified, and since actin is likely to be required for EPS secretion^24,25^, an alternative model where force is generated from EPS polymerization has not been excluded^15,16^.

Within each photosynthetic lineage, loss of photosynthesis led to secondary heterotrophs, many of which are parasites that derive energy from their host^26^. Transitions to epizoic^27^ and free-living^28,29^ heterotrophy are also well-documented. However, in most of these cases, the manner of energy acquisition remains unclear. In the diatom genus *Nitzschia*, loss of photosynthesis led to a group of free-living heterotrophs^30,31^. These apochlorotic diatoms have been isolated from the nutrient-rich waters of the intertidal zone where they occur as epiphytes on seaweeds, on decaying plant matter, and in the surrounding waters^30,32–35^. As with many photoautotrophs that transition to heterotrophy^28^, they have retained their plastid genomes and certain plastid-localized metabolic functions, but have lost key photosynthetic genes^36^. Early work showed that apochlorotic diatoms can grow on a variety of simple and complex carbohydrates including cellulose and the red algal cell wall polysaccharides agarose and carrageenan^32,33,37^. Recent genome sequences identify lignin-degrading enzymes in *Nitzschia* Nitz4^38^ and expansion of secreted proteins and functions related to organic carbon acquisition in *Nitzschia putrida*^39^. Thus, candidates for key heterotrophy-related functions are beginning to emerge.

Here, we isolate new strains of apochlorotic diatoms from Singapore’s intertidal zone. Live-cell imaging documents EPS trail deposition and complex motility-related behaviours that include high force generation (~800 pN), cooperative motility and collision-induced reversal. Variations in motility and metabolism suggest that apochlorotic diatoms are undergoing substantial ecophysiological radiation. We propose that these new isolates provide excellent models to study the evolutionary transition to free-living heterotrophy.

## Results

### Isolation and characterization of apochlorotic diatoms

Diatoms were cloned from organic materials collected from the intertidal zone on Sentosa island, Singapore (see Materials and Methods). Five clones were initially isolated from decaying plant matter, the brown alga *Sargassum*, and the green alga *Bryopsis* (Fig. 1A). Subsequent work revealed the ability of these diatoms to metabolize the brown algal cell wall polysaccharide alginate (see below). Thus, we isolated an additional seven clones from *Sargassum*. Phylogenetic analysis indicates that these 12 isolates fall into three distinct clades (Fig. 1A). Isolates were named *Nitzschia* singX-Y, where X designates the clade number and Y, the isolate number. Isolates in clade 1 and 2 are sister taxa, with clade 2 having an affinity for *N. alba*, while clade 3 is distantly related to clades 1 and 2.

**Figure 1:**
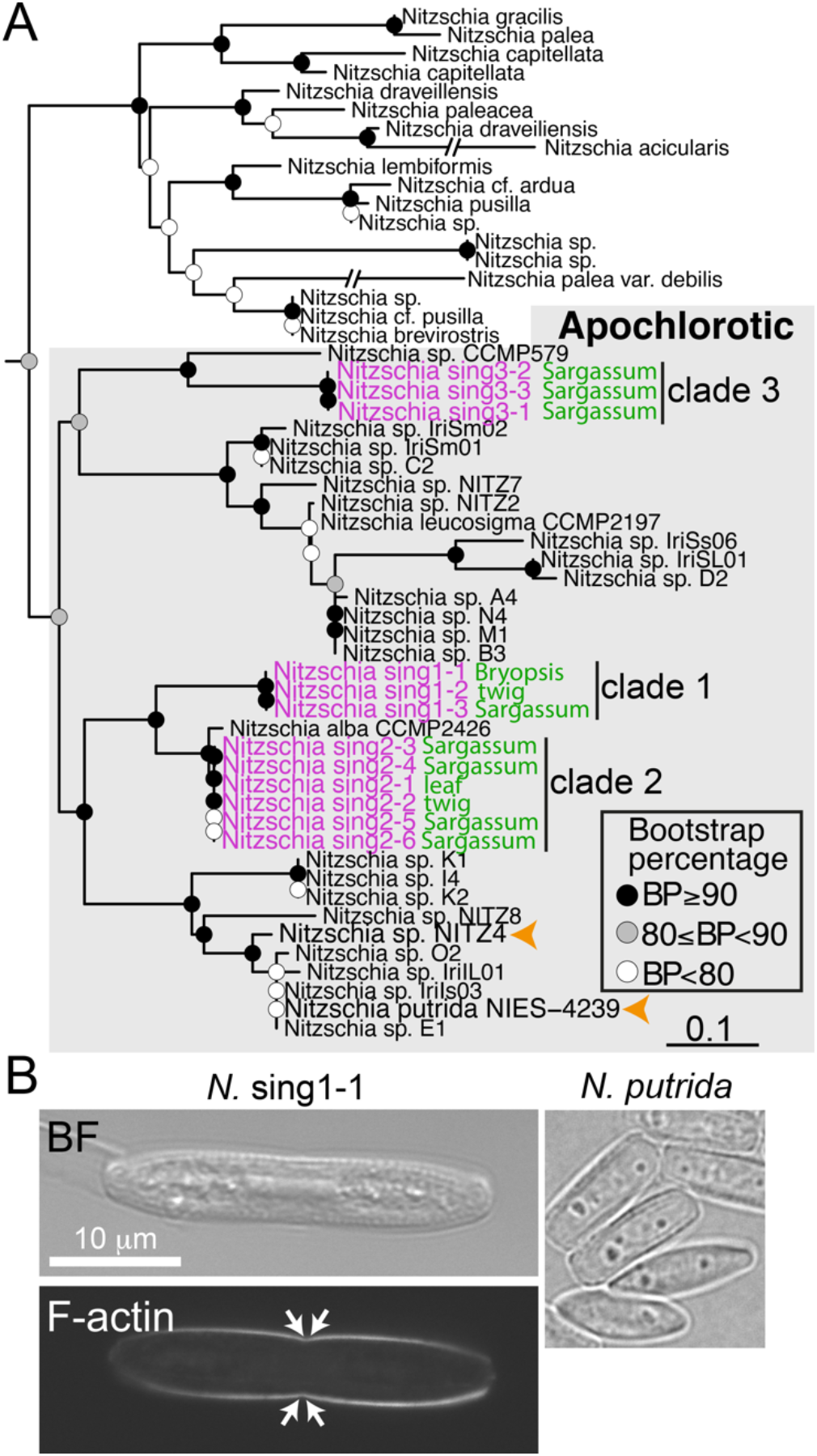
Characterization of Singaporean apochlorotic diatoms. **(A)** Maximum likelihood phylogeny of apochlorotic *Nitzschia* (shaded box) and Bacillariales photosynthetic outgroup taxa. The Singaporean isolates are identified in magenta. The material from which they were isolated is indicated in green. Filled circles show simplified bootstrap support values. Scale bar = 0.1 nucleotide substitutions. Orange arrow-heads identify apochlorotic species with sequenced genomes. **(B)** Factin staining of *N.* sing1-1. The arrows point to the approximate position of proximal raphe ends. A bright-field (BF) image of *N. putrida* is shown for comparison. Scale bar = 10 μm.

Isolated diatoms grown on agarose seawater^40^ media form a radially expanding colony. Growth rates vary substantially both within and between clades. Rates of colony expansion for clade 1 and 2 diatoms vary between 100 and 300 nm/s (Fig S1A) By contrast, all clade 3 diatoms and the recently sequenced apochlorotic *N. putrida*^39^ show very little colony expansion, with cells dividing to form aggregates at the site of inoculation (Fig. S1B and C).

Clade 1 diatoms were isolated from green and brown algae, and decaying plant matter, suggesting that they occupy diverse ecological niches. They are also among the fastest growing isolates. Thus, we chose *N.* sing1-1 as our model apochlorotic diatom (hereafter referred to as *N.* sing1). F-actin staining reveals characteristic bands underlying the raphe (Fig. 1B), and scanning electron microscopy (SEM) of frustules identifies eccentric raphes, hymenate pore occlusions and strongly hooked distal raphe ends (Fig. S1D).

#### Growth on algal polysaccharides and catabolite repression

We next examined the growth of *N.* sing1 on seawater media solidified with red algal cell wall polymers agarose and carrageenan. As previously observed with *N. alba*^37^, both of these substrates could be utilized as the sole carbon source (Fig. S2). In addition, *N.* sing1 grows on the brown algal polysaccharide alginate. For each polysaccharide substrate, the rate of radial colony expansion generally has a concentration optimum and tends to decrease with increasing concentration (Fig. S2). In the case of alginate, the medium underwent liquefaction and browning indicative of polysaccharide hydrolysis (Fig. 2A). This is confirmed by a heat-sensitive alginate lyase enzyme activity detected in the media of *N.* sing1 grown with alginate but not glucose (Fig. 2B). Representatives of each clade of Singaporean diatoms liquefy alginate, but *N. putrida* does not. This suggests that modes of heterotrophy vary substantially within the apochlorotic lineage. Neither agarose nor carrageenan undergo liquefaction. However, on these media the diatoms tunnel to grow invasively (Fig. S3) as has been documented for *N. albicostalis*^33^ and *N. alba*^37^.

**Figure 2:**
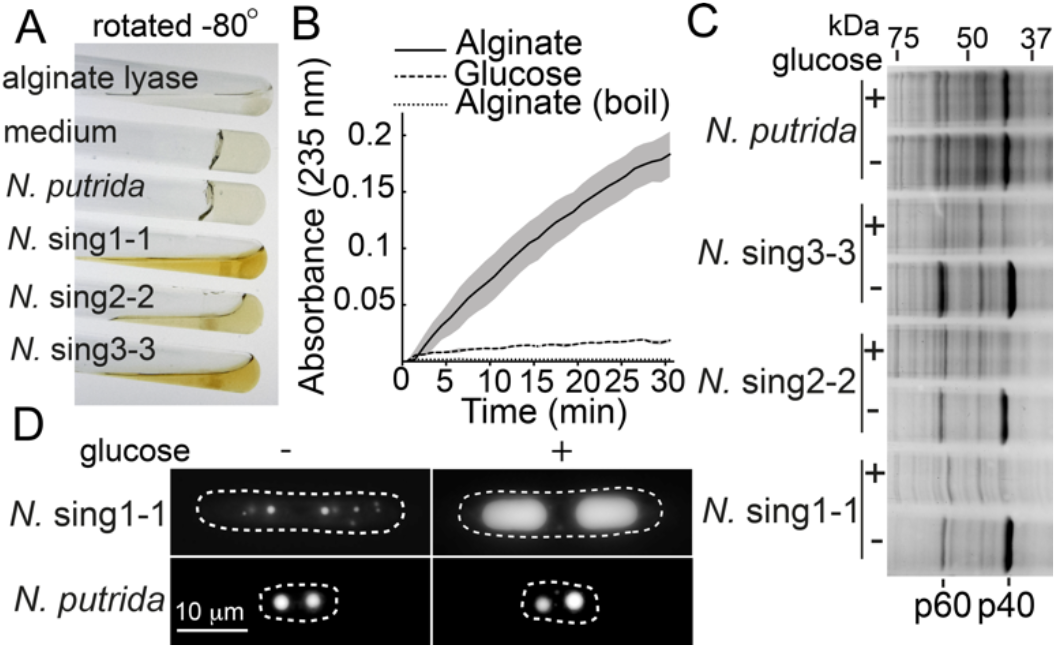
Diatom alginate utilization and catabolite repression. **(A)** The image shows cultures of diatoms on a 1.5% seawater alginate medium. The tubes are rotated approximately −80 degrees to show alginate gel liquefaction. Alginate lyase enzyme and medium alone serve as positive and negative controls, respectively. **(B)** The graph shows *N.* sing1-1 alginate lyase enzyme activity in the indicated media after 4 days of growth. Note that boiling (boil) abolishes the enzyme activity. **(C)** SDS-PAGE of total cell extracts from diatoms grown on seawater agar medium in the absence (-) or presence (+) of 0.5% glucose. Glucose-repressed proteins species p40 and p60 are identified. **(D)** Lipid droplet accumulation of *N.* sing1-1 and *N. putrida* on seawater agarose media in the presence (+) and absence (-) of glucose. The dotted line traces the cell periphery. Scale bar = 10 μm. Related to Figure S5.

We next examined cellular extracts of diatoms grown on seawater agar medium with and without 0.5% glucose. SDS-PAGE reveals two *N.* sing1 proteins, p40 and p60, that are abundant on agarose media, but substantially diminished when glucose is present. This repression is observed for representatives of each *N.* sing clade, but not *N. putrida* (Fig. 2C). Alginate also promotes catabolite repression in *N.* sing1, indicating that it is also a preferred carbon source (Fig. S4A). *N.* sing1 accumulates large lipid droplets in the presence but not absence of glucose. By contrast, *N. putrida* lipid droplets have a similar appearance irrespective of glucose presence (Fig. 2D and S4B). This provides further evidence for the metabolic responsiveness of *N.* sing1 to a preferred carbon source.

#### Environmental control of EPS trails and motility

While measuring *N.* sing1 growth, we found that the EPS can be seen as a refractive trail by bright-field microscopy (Fig. 3A). This is likely due to swelling of the EPS to form a refractive convex cross-sectional profile. EPS trails formed on 1% agarose have a uniform width and appearance. By contrast, on 2% agarose, where motility is substantially diminished, the trails take on a broken appearance and the EPS forms refractive spherical structures. To examine how the availability of seawater affects motility, we overlayed the medium with seawater (Fig. 3B). In this condition, a dramatic increase in the speed of motility is observed as compared to plates without a seawater overlay (Fig. 3C and D). Here, trails are not seen because the EPS is not at the air interface. Together, these findings link EPS condensation behaviour with motility, and suggest that nascent EPS function is sensitive to timely hydration. Interestingly, diatoms are also observed gliding in a monolayer at the seawater-air interface, indicating that *N.* sing1 motility is not strictly dependent on substratum attachment (Fig. 3C).

**Figure 3:**
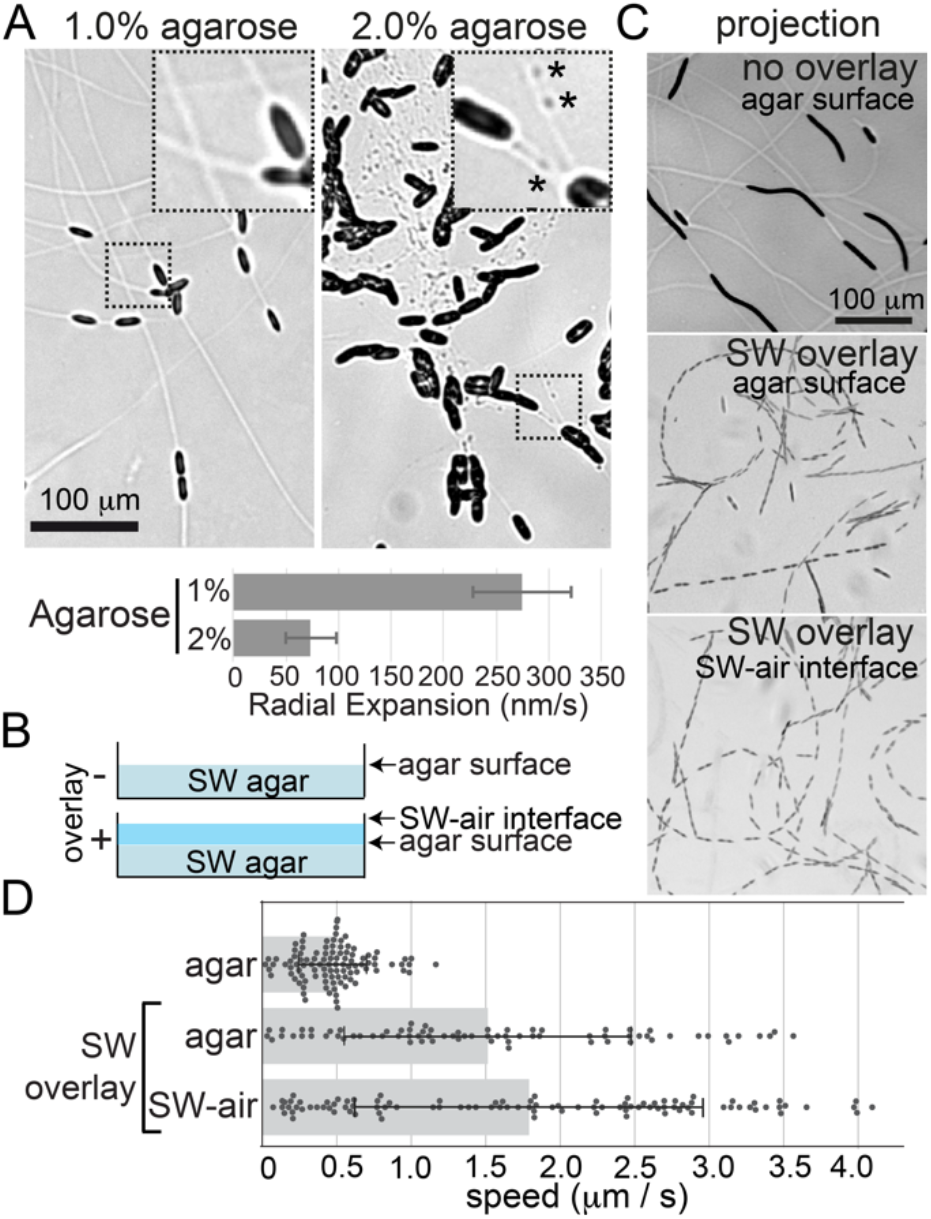
Environmental control of EPS and motility. **(A)** Appearance of EPS trails on 1 and 2% seawater agarose medium. The dashed box is magnified on the upper right. Scale bar = 100 μm. Asterisks identify refractive EPS puncta. The graph shows the speed of growth on each medium. **(B)** The diagram depicts a side view of assay plates in the presence (+) or absence (-) of a seawater (SW) overlay. **(C)** The projections show diatom motility under the indicated conditions. Total time = 305 seconds. Scale bar = 100 μm. **(D)** The graph shows the speed of diatoms shown in (C). The bar shows the average with standard deviation indicated. Related to Supplementary Movies 1, 2 and 3.

#### Cooperative motility, force generation, and force sensing

Movies of EPS deposition allowed us to differentiate diatoms laying fresh EPS trails from those moving on pre-existing trails (Fig 4A). We refer to these as trail blazers (blazers) and trail runners (runners), respectively. Blazers instantly accelerate upon joining a trial, while runners that leave a trail instantly decelerate (Fig. 4B to E). These observations indicate that gliding motility is inherently more efficient when occurring on an EPS trail. Because of this relationship, runners tend to catch up to blazers to form chains of cells, particularly at the colony’s expanding edge. When fast-moving runners catch up to blazers, the blazer can instantly accelerate (Fig. 4F and G). Thus, runners can exert pushing forces to make blazers move faster.

**Figure 4:**
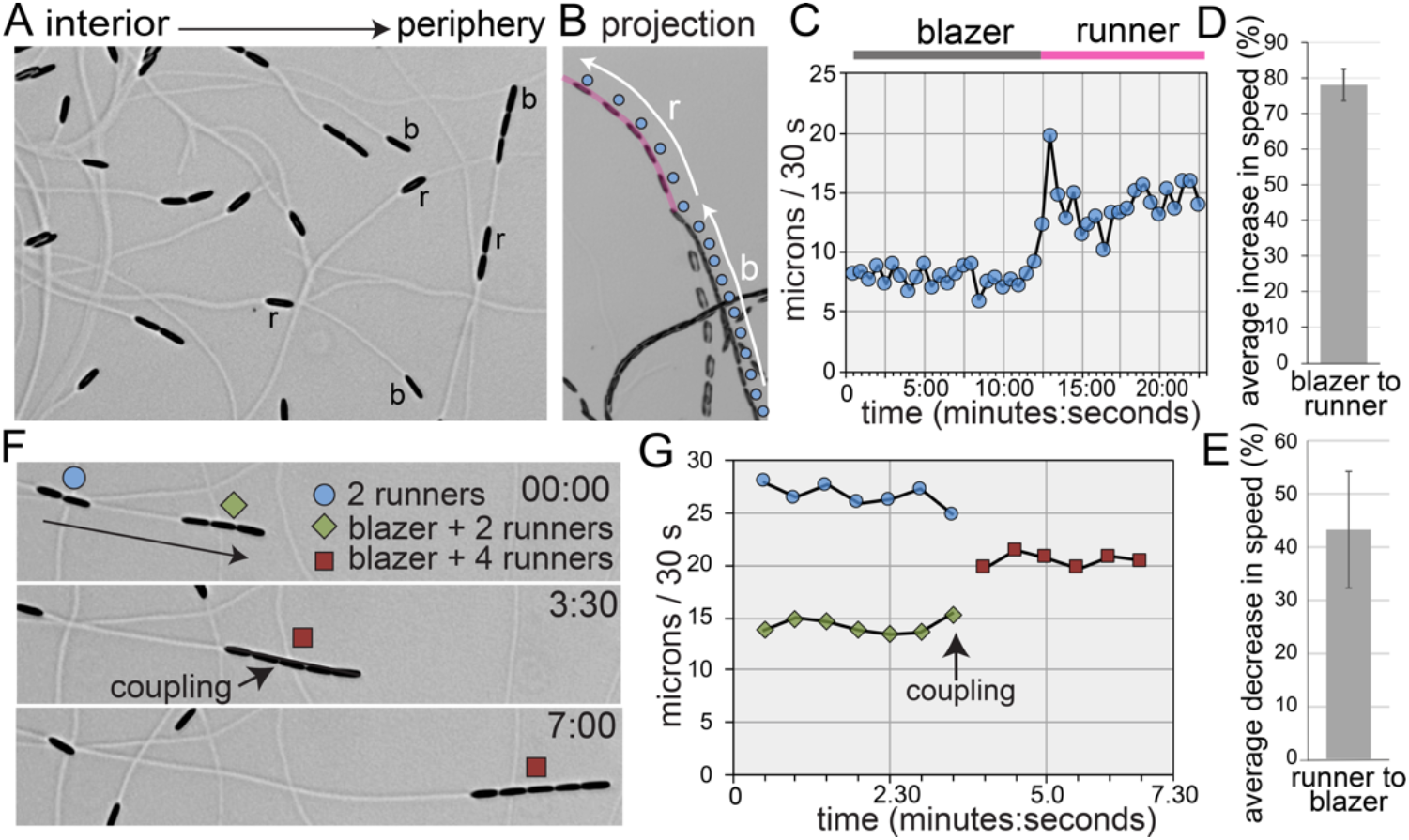
Live-cell imaging of gliding and EPS deposition by *N.* sing1-1. **(A)** The image shows a field of diatoms near the colony edge. The direction from interior to colony periphery is indicated. Diatoms laying a new trail are blazers (b), while diatoms gliding on a pre-existing trail are runners (r). Note that these identities are interchanged as diatoms reverse or transition on or off trails. Related to Supplementary Movie 4. **(B)** A blazer (blue circle) instantaneously accelerates upon joining a pre-existing trail (magenta line). White arrows indicate the periods spent as blazer (b) and as runner (r). The projection is derived from Supplementary Movie 5. **(C)** The graph shows the speed of the diatom from (B). **(D)** The graph quantifies the average increase in speed when a blazer becomes a runner. Standard deviation is indicated (n=6). **(E)** The graph quantifies the average decrease in speed when a runner becomes a blazer. Standard deviation is indicated (n=6). **(F)** Runners can push blazers to make them move faster. A chain consisting of a blazer and two runners is caught by a pair of runners to make a chain of five. The groups are identified at the indicated frames by symbols given in the legend. The moment when cell-cell coupling occurs is indicated. Related to Supplementary Movie 6. **(G)** The graph shows the speed of the groups as defined in (F).

Mathematical modeling shows that the movement of diatoms is well approximated by Brownian motion over large space and time scales (Supplementary information). A quasi-steady state analysis of the model provides a mathematical relationship between motion characteristics and establishes that colony diffusivity increases with diatom speed. This relationship is preserved in runner-blazer groups, which exhibit an increase in speed compared to lone blazers (Fig. 4F and G). Thus, modeling confirms the tendency for cooperative motility to increase colony diffusivity.

Broadside collisions between diatoms are readily observed and these frequently lead to reversal of the impacting diatom. In Figure 5A, diatom 1 undergoes three successive impacts with a relatively stationary diatom 2. The initial collision is followed by a rapid reversal. However, the second and third collisions are characterized by longer periods spent pushing. This is coincident with increasing degrees of deflection of diatom 2 (Fig 5A). Thus, the capacity for force generation appears to be dynamic. Not all collisions lead to reversal. In some cases the impacting diatom slows dramatically upon collision and continues to move forward as it pushes the other diatom out of its path (Fig 5B and C). Together, these observations reveal the ability of moving *N.* sing1 diatoms to impart force, which impacted diatoms resist through substratum adhesion.

**Figure 5:**
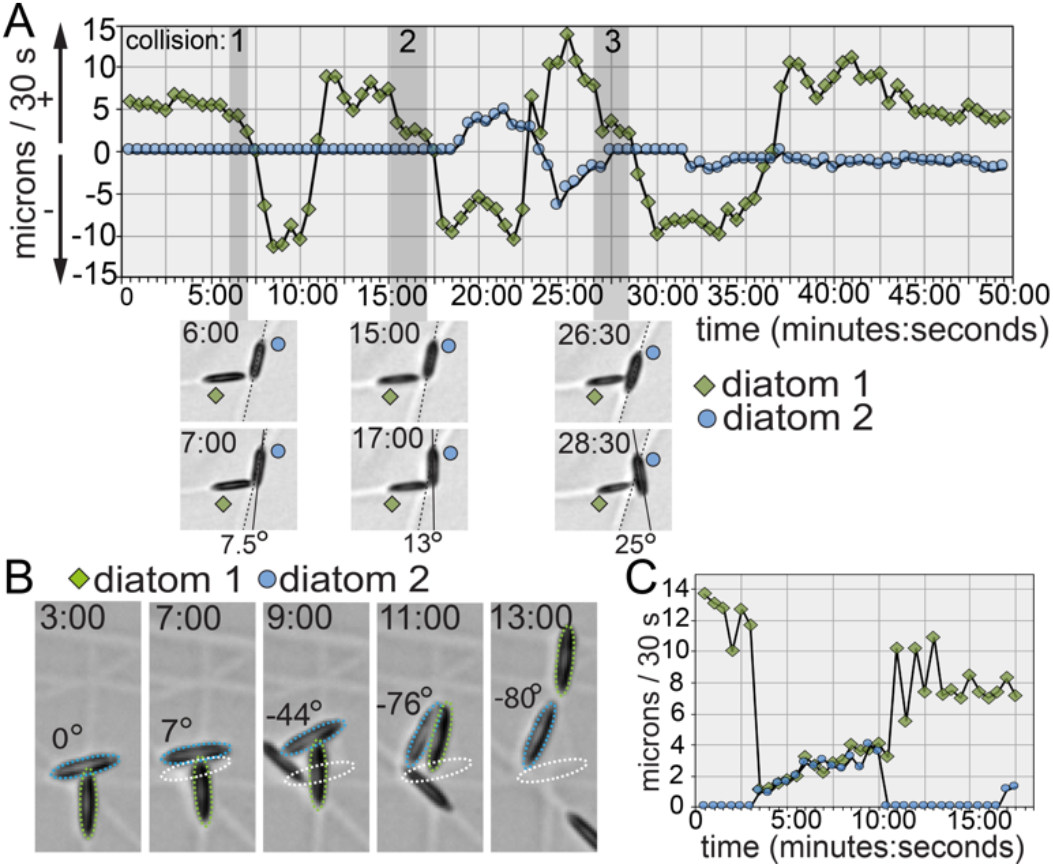
Collision-triggered reversal and pushing. **(A)** Diatom 1 collides with a stationary diatom 2 three consecutive times. Frame-to-frame speed is shown in the graph. The images at the bottom show the first (upper panel) and last (lower panel) frames of the collisions. Related to Supplementary Movie 7. **(B)** Diatom 1 collides into a stationary diatom 2, which is pushed laterally and rotates until diatom 1 frees itself. The original position of diatom 2 is given by the dotted white outline and its degree of rotation is indicated numerically. Related to Supplementary Movie 8. **(C)** The graph shows the speeds of diatom 1 and diatom 2 from (B).

We next employed a method that allows the measurement of forces exerted by single diatoms. Here, a force-calibrated glass micropipette is placed in the path of moving diatoms and force is estimated through the degree of pipette deflection^41,42^. These data reveal forces between ~100 and ~800 pN (Fig. 6A, B and S5). Lower force measurements are associated with glancing contact with the pipette. By contrast, high force measurements occur within the context of head-on contact and adhesion between the diatom and pipette. Adhesion is evidenced by diatom detachment from the agar surface at a load of ~740 pN (Fig. 6B #3) and an apparent EPS tether that attained a length of ~3-4 μm before snapping at a load of ~800 pN (Fig. 6B #2). Diatoms pass underneath the pipette while it undergoes deflection suggesting that they are subject to downward pushing forces of ~84 pN while pushing/pulling on the pipette (based on an estimated diatom height of 4 μm). Overall, these data show that *N.* sing-1 is capable of producing surprisingly high forces. For comparison, single intact muscle filaments produce forces of ~200-300 pN^43,44^.

**Figure 6:**
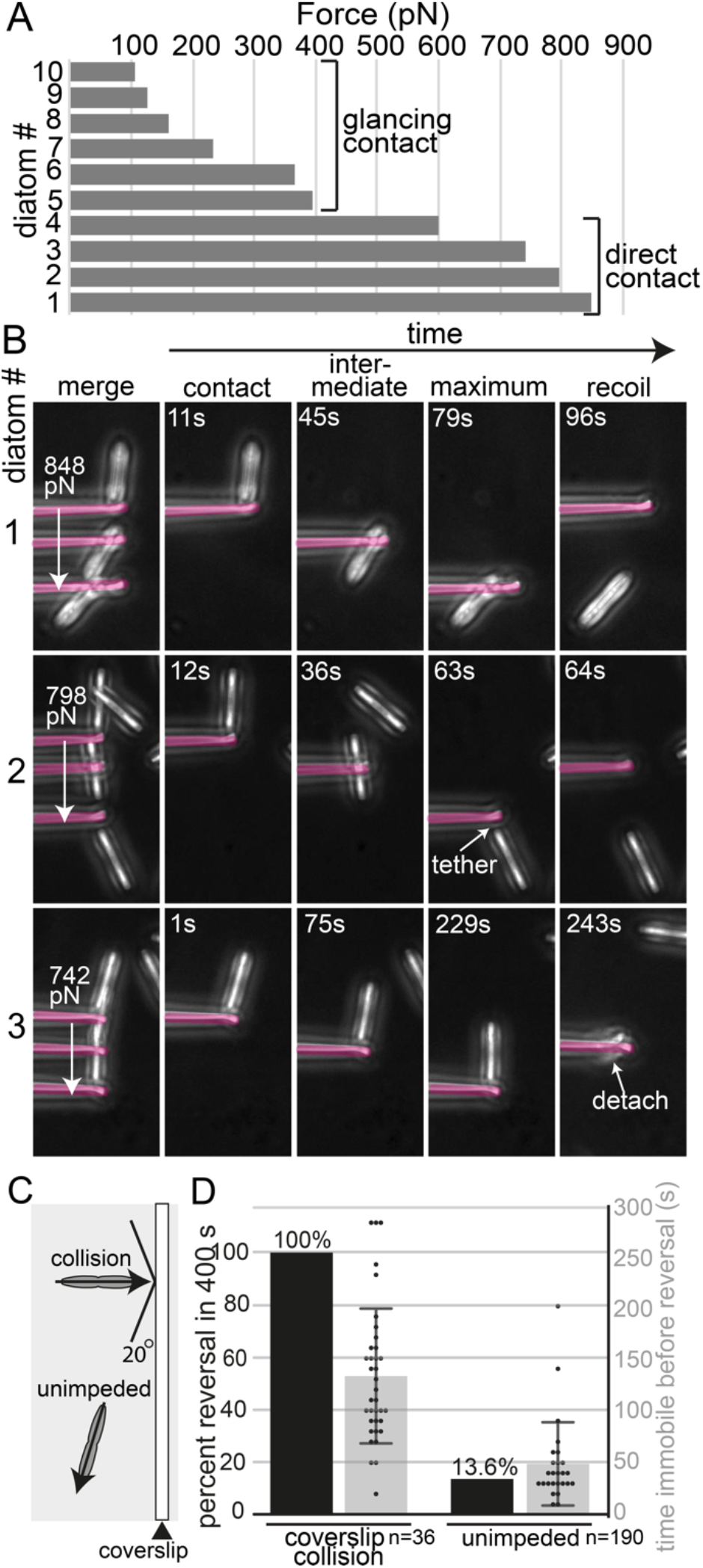
Forces exerted by *N.* sing1 diatoms and mechanosensing. **(A)** The graph shows the maximum force produced by single diatoms pushing/pulling on the pipette. **(B)** Images taken from the indicated movies show first contact (contact), an intermediate timepoint (intermediate), the point of maximum pipette deflection (maximum) and pipette recoil (recoil). The arrows in the merge panel show maximum force values. The pipette is overlayed in opaque magenta. Note that for measurements 1 and 2 the diatom passes underneath the pipette. Related to supplementary movies 9, 10 and 11. **(C)** The schematic shows the experimental set-up for immobilization-triggered reversal. Only diatoms that hit the wall at an angle of >20 degrees are included in the dataset. **(D)** The graph shows the percentage of cells reversing within 400 seconds (left y-axis). The scatterplot (right axis) shows the duration of time spent immobile prior to reversal. The light grey bar identifies the mean. Standard deviation is indicated.

Collision-associated reversals suggest that *N.* sing1 diatoms can sense force. However, reversals do not always follow collisions (Fig. 5C and D) and also occur in free-running diatoms. To isolate collision events, we grew diatoms into a coverslip embedded perpendicular to the agar surface and compared these diatoms to nearby free-running diatoms that did not experience collision (Fig. 6C). In this experiment, 100% of colliding diatoms reverse within 400 seconds of being immobilized. By contrast, 13% of individuals whose motility is unimpeded reverse within the same time interval (Fig. 6D). In these experiments, colliding diatoms spend a longer period immobile prior to reversal as compared to free-running reversals. Free-running diatoms also slow prior to stopping and reversing, suggesting underlying distinctions between free-running and collision-induced reversals (Fig. S6). Irrespective of this, these data show that collisions that impede forward motion significantly increase the probability of reversal. This type of force sensing is likely to increase colony diffusivity especially where substrates possess complex morphologies.

## Discussion

The apochlorotic diatoms described here were isolated from a variety of organic materials (Fig. 1A) and grow on a broad range of algal polysaccharides (Fig. 2A, and S2). Evidence for catabolite repression (Fig. 2C and S4A) identifies an aspect of metabolism classically associated with heterotrophy in fungi^45^ and bacteria^46^. Overall, these findings are consistent with a general role for apochlorotic diatoms in coastal marine nutrient cycling—one akin to the role of osmotrophic fungi^47^ in terrestrial environments. Interestingly, unlike the diatoms identified here, *N. putrida* does not metabolize alginate nor show evidence for catabolite repression (Fig. 2A and C). A high degree of ecophysiological variation is further evidenced by poor motility of clade 3 isolates and *N. putrida* as compared to clade 1 and 2 isolates (Fig. S1A to C). Together, these findings suggest that apochlorotic diatoms exploit distinct feeding strategies and are undergoing substantial evolutionary radiation.

A series of evolutionary innovations culminating in force generation from motility are likely to have predisposed the diatoms to a successful transition to heterotrophy. These include the advent of the silica-based cell wall, bilateral symmetry, the raphe, and forceful motility. Certain marine bacteria are highly evolved for alginate metabolism^48,49^, but are unlikely to generate forces necessary for tunnelling^50^. Thus, high forces from diatom gliding motility (Fig. 6A and B) are likely to underlie invasive growth (Fig. S3) and allow access to nutrient pools unavailable to competing microorganisms. This is consistent with *N. alba* invasive growth on brown algal tissues^37^, and tunneling in both *N. alba* and *N. albicostalis*, which appears to be stimulated by the presence of bacteria^33,37^. In terrestrial environments, the fungi have a similar advantage where force from pressurized hyphal networks underlies invasive growth^51^. Thus, distinct manners of force generation appear to provide an advantage to eukaryotic heterotrophs over their bacterial counterparts.

Diatom EPS trails have been visualized and characterized by electron microscopy^52^, atomic force microscopy (AFM)^53,54^ and various staining techniques^15,17,19,55^. To the best of our knowledge, this is the first study to observe EPS deposition by bright field microscopy. On agarose concentrations that favour motility, uniform refractive EPS trails are presumably visible due to their convex cross-sectional profile. With increasing agarose concentration, motility slows substantially and EPS trails lose their uniform cross-sectional profile to form aberrant refractive puncta (Fig. 3A). This suggests that freshly secreted EPS undergoes rapid swelling that is sensitive to timely hydration and/or availability of critical seawater ionic constituents. A critical role for hydration is also consistent with dramatically enhanced motility when agar plates are overlayed with seawater (Fig. 3B to D). The sea surface microlayer (SSM) is known to have a distinct physical, chemical, and biological composition^56^. The finding that gliding occurs in a monolayer at the SSM (Fig. 3C) suggests that the EPS has an affinity for the seawater surface underside. This provides a physical basis for previous work showing that apochlorotic diatoms are enriched at the SSM^34^.

The relationship between runners and blazers leads to cooperative motility (Fig. 4), and is likely to be related to the dual function of EPS in adhesion and motility. Runners that go off-trail instantly decelerate. This is consistent with more nascent EPS being consumed by the adhesive function. By contrast, blazers joining a trail instantly accelerate (Fig. 4B-D) because adhesive contacts are already present, and only EPS-EPS contacts are required. Interestingly, line scans of trails do not change dramatically between a fresh trail and one that has been passed over by additional runner diatoms (Fig. S6). Thus, runners may be secreting substantially less EPS than blazers. This could also factor into their tendency to move at higher speeds.

The available data on diatom motility do not distinguish between force generation through actomyosin motors and EPS polymerization^15,16^. Our force measurements tend to support the former model. Single myosin molecules exert forces of 3-4 pN^57^, while isolated muscle filaments can generate maximum forces of ~200-300 pN^43,44^. Thus, *N.* sing1 peak forces of 700-800 pN (Fig. 6A and B) are consistent with the cooperative action of multiple myosin motors arrayed along the length of raphe-associated actin filaments. Force measurements were made with single diatoms and are likely to be higher in runner/blazer chains. Thus, cooperative motility is expected to increase both colony diffusivity (Supplementary Information) and the ability to tunnel effectively.

A related set of equations describe colony diffusivity of *N.* sing1 (Supplementary information) and the mixotrophic diatom *Navicula*^58^. Thus, periodic reversal combined with random turning is likely to be a general strategy used by raphid pennate diatoms to avoid immobilization after encountering an obstacle. By sensing immobilization (Fig. 6C), *N.* sing1 is able to reduce the period of immobility, while presumably maintaining an independent frequency of free-running reversal. Distinct mechanisms could underlie *N.* sing1 force sensing. For example, mechanosensitive ion channels are established force sensors that could trigger a signalling cascade leading to reversal. In another model, strain on the force-generation machinery could trigger reversal through a feedback mechanism. More work will be required to determine the mechanism of force sensing.

The transition from photoautotrophy to obligate heterotrophy is likely to have been accompanied by a variety of physiological adaptations. However, because many pennate diatoms are higly evolved mixotrophs^59–62^, it remains unclear whether alginate utilization, cooperative motility, and force sensing are unique to apochlorotic diatoms or predate their emergence. Identifying the genetic basis for diatom obligate heterotrophy will require an integrated approach that combines comparative genomics with molecular, biochemical, cellular and physiological studies.

## Acknowledgements

We thank Judy Lim for help with making movies, Aditi Lakshmanan and Jie Yun Wong for technical assistance, and Matt P. Ashworth at the UTEX Culture Collection of Algae at the University of Texas, Austin for assistance with SEM documentation. GJ thanks the National Parks Board, Singapore for permission to collect, and Danwei Huang for helpful information on Sargassum biology and distribution. Work in the Jedd Group is supported by the Temasek Life Sciences Laboratory. AA acknowledges funding from the National Science Foundation (DEB 1651087). JMN and YA are supported by the Natural Sciences and Engineering Research Council of Canada (NSERC: RGPIN-2019-06435, RGPAS-2019-00014, DGECR-2019-00321). KK and YJ acknowledge support from the Singapore Ministry of Education, Academic Research Fund Tier 2 (MOE-T2EP50220-0015), and the National Research Foundation, Singapore, and the Ministry of Education under the Research Centres of Excellence program. This paper was typeset with the bioRxiv word template by @Chrelli: www.github.com/chrelli/bioRxiv-word-template

## Author contributions

GJ conceived the project. PZ, CQ, ZHL, ZTY, AA, KK, and GJ conducted experiments. PZ, CQ, ZHL, ZTY, AA, KK, YJ and GJ analysed the data. YA and JN conduced mathematical analysis contained in Supplementary Information. GJ wrote the paper with inputs from all the co-authors.

## Competing interest statement

The authors have no competing financial interests.

## Materials and Methods

### Diatom isolation

Organic materials were collected at low tide from the intertidal zone on Sentosa island, Singapore (latitude 1.259895, longitude 103.810843; Singapore National Parks Board permit #NP/RP20-016). Samples were inoculated at the centre of synthetic seawater^40^, 2 % (w/v) agar plates supplemented with 100 μg/ml ampicillin (Sigma-Aldrich, A9518) and 50 μg/ml kanamycin (Sigma-Aldrich. K4000). After two to three days of incubation at 30°C, single diatoms gliding away from the source inoculum were excised with a scalpel and transferred to a fresh plate. To ensure that all isolates are single cell clones, the isolation process was repeated. Diatom clones were cryopreserved according to the method of Stock *et al.*^63^. The *Sargassum* species present at the collection site were identified as a mixture of *S. polycystum, S. cf. granuliferum* and *S. ilicifolium* (Fig. S8) through a maximum likelihood (ML) analysis of ITS-2 alignment as previously described^64^. New sequences are available in GenBank under accessions OQ165106 to OQ165109. *N. putrida* (strain #NIES-4239) was obtained from The Microbial Culture Collection at the National Institute for Environmental Studies 16-2, Onogawa, Tsukuba, Ibaraki 305-8506, Japan.

### Diatom Phylogenetic Analysis

We sequenced one nuclear gene (28S d1-d2 rDNA) and two mitochondrial genes *(cob* and *nad1)* for each of the Singapore isolates following the PCR and sequencing protocols outlined in Onyshchenko *et al*.^31^. Raw Sanger sequences were edited and assembled with Geneious ver. 7.1.4 (Biomatters Ltd.). New sequences are available in GenBank under accessions OQ319050-OQ319073 and OQ317929-OQ317940. The proteincoding *cob* and *nadl* genes were aligned by eye in Mesquite ver. 3.6^65^ using the predicted amino acid translations as a guide to preserve codon structure. Partial 28S rDNA genes were aligned using SSU-ALIGN ver. 0.1^66^ with a heterokont-specific covariance model^67^. The three alignments were concatenated with AMAS^68^. We used IQ-TREE ver. 1.6.12^69^ for phylogenetic reconstruction. The concatenated alignment was partitioned by gene and codon position, and IQ-TREE’s partitioning and model selection procedure (-m TESTMERGE) was used to identify the best fitting partition and nucleotide substitution model^70,71^. The model included used edge-proportional branch lengths (IQTREE’s ‘-spp’ option) to account for differences in evolutionary rates among partitions. Branch support was based on 1,000 ultrafast bootstrap replicates^72^.

### Diatom growth, measurement, microscopy, and movies

Diatoms were cultured on a synthetic seawater medium^40^ with the following modification. NaCl was employed at a final concentration of 180 mM, and for growth in liquid cultures Na_2_SiO_3_ was employed at a final concentration of 840 μM. For solid media, salt solutions I and II were prepared as 4X stock solutions, while polysaccharides were prepared at 2X concentrations. Where necessary, polysaccharides were boiled, and equilibrated to 60°C before mixing with salt solutions I and II to yield 1X final concentrations. Polysaccharides were obtained from the following sources: agar (BD, 214010), agarose (Vivantis, PC0701), carrageenan (Sigma-Aldrich, C1013), low viscosity sodium alginate (Sigma-Aldrich, A1112) and medium viscosity sodium alginate (Sigma-Aldrich, A2033). For the alginate liquefaction assay (Fig. 2C) medium viscosity alginic acid was employed.

To measure the speed of radial colony expansion (Fig. S1A and S2), a starter culture was prepared on a seawater agar petri dish. From these confluent plates, a small block (~0.5 X 0.5 cm) was excised with a scalpel and used to inoculate fresh plates from which measurements were derived. After overnight growth at 30°C, the front of the radially expanding colony was marked and then marked again after an additional 24 hours. These marks were used to calculate speed, in nm/s. Each speed measurement was done in triplicate.

For movies, diatoms were grown on seawater agar medium in petri dishes. Movies were made using an Olympus BX51 upright microscope equipped with a 5x objective using a Photometrics CoolSNAP HQ (Teledyne Photometrics) camera controlled by MetaVue software (Molecular Devices). Frames were acquired every 30 seconds at an exposure time of 20 milliseconds. For movies shown in Figure 3C frames were acquired every 5 seconds. Graphs of diatom speed were made by manually measuring frame-to-frame diatom movement in ImageJ (https://imagej.nih.gov/ij/index.html). For the quantification shown in Figure 4D and E, the average increase or decrease in speed was calculated from the average speed over 6 frames before and after joining the trail.

To examine total cell proteins(Fig 2C and Fig S4A) by sodium dodecyl sulfate-polyacrylamide gel electrophoresis (SDS-PAGE), plates on which diatoms had been grown to confluence were flooded with 7.5 ml phosphate buffered saline+ (PBS+) (10 mM Na_2_HPO_4_, 1.8 mM KH_2_PO_4_, 240 mM NaCl) and cells were scraped off gently using a cell scraper (Corning Incorporated Costar, 3010). The cells were pelleted by centrifugation at 3.9 k x *g* for 10 minutes and washed once with 7 ml of PBS+, before being lysed by boiling in SDS-PAGE loading buffer. Proteins were resovled using 10% polyacrylamide gels.

For staining with phalloidin (ThermoFisher, A12379) and DAPI (ThermoFisher, D1306) (Fig. 1B), a block of seawater agar medium from a confluent plate was transferred to the center of a coverslip. The coverslip was placed in a 60 mm petri dish, overlayed with 6 ml of liquid synthetic seawater medium (0.5% (w/v) glucose), and incubated at 30°C for 24 hours. The agar black was removed, and the coverslip placed cells-up on a piece of parafilm in a 90 mm petri dish. The cells were washed once with PBS+ and then fixed in PBS containing 4% paraformaldehyde (Electron Microscopy Sciences, 15713) for 30 minutes at room temperature. Samples were washed 3 times with PBS+ and then incubated with 0.17 μM phalloidin 488 (stock solution: 66 μM in DMSO) in PBS+ for 1 hour in the dark. Samples were washed with PBS+ (3x fast wash + 3x 10 minutes wash) at room temperature. After the last wash, 5 μl mounting medium (PBS, 20% glycerol, 2 μg/ml DAPI, 1X antifade), (100x antifade stock: 20%(w/v) *n*-propyl gallate (Sigma P3130) in dimethyl sulfoxide) was added and the slip was placed on a microscope slide cells down and sealed with parafilm. For staining of lipid droplets with BODIPY, diatoms were grown on seawater agar media. After 2 days growth at 30°C, a block (~1 cm x 1 cm) was excised with a scalpel and transferred cells-up onto a microscope slide. The diatoms were overlayed with 4 μl of a solution containing 30 μg/ml BODIPY™ 505/515 (Invitrogen, D3921) in PBS+. After approximately 30 minutes a coverslip was added and the diatoms were imaged by fluorescence microscopy. The diatoms frustules were prepared for scanning electron microscopy (SEM) as previously described^73^.

### Force measurement

To measure the force produced by diatom movement (Fig. 6A and B), a small block, approximately 0.25 x 0.5 cm, was excised from a starter culture and used to inoculate 1.5% agarose seawater medium contained by a u-shaped thin-wall chamber made of polydimethylsiloxane (DOW, 4019862) on top of a coverslip^41^. After incubation at room temperature for one day, the chamber was flooded with seawater (Electrostatic attraction between the pipette and medium necessitated that these experiments be conducted with a seawater overlay as in Fig. 3B). The chamber was then placed on a microscope stage with a slide holder and a micropipette, prepared as described below, was inserted through the open side of the chamber and positioned in front of a diatom using an MP-285 motorized micromanipulator (Sutter Instruments). Movies were made using an Olympus IX81 equipped with a 40x objective manipulated with MetaMorph software (Molecular Devices), with frames acquired every 0.5 seconds. Frame-to-frame micropipette and diatom movement were manually measured using ImageJ, and used to produce force/velocity graphs (Fig. S5).

Micropipette production: Briefly, thin wall glass capillaries (1 mm outer diameter, 0.78 mm inner diameter; World Precision Instruments, TW100F-6) were pulled using a P-97 micropipette puller (Sutter Instruments) and the tip was cut to the desired inner diameter of approximately 2 μm with an in-house developed heated platinum wire. The stiffness of the micropipette was calibrated using a standard micro glass rod (*k_s_* = 21.09 ± 4.22 pN/μm). The preparation of standard micro glass rod the calibration of the working micropipette were conducted as previously described^41^. Before use, the tip of the micropipette was incubated overnight with 3% fetal bovine serum (Sigma-Aldrich, A7030) and incubated overnight.

### Alginate lyase enzyme activity assay

Diatoms were grown in liquid medium for 3 days at 30°C, after which cells were removed using a cell strainer (SPL Life Sciences, 93040). The medium was then centrifuged at 3.9 k x *g* for 15 minutes and concentrated using a Pierce™ protein concentrator with a 3 kilodalton cut-off (ThermoFisher, 88526). The concentrated medium was diluted 1:5 with synthetic seawater before use. For heat-inactivated controls, the diluted culture media was heated at 100°C for 5 minutes and then briefly centrifuged. To perform the enzyme assay, 5 μL of diluted sample was added to 45 μL sodium alginate buffer (10 mM Tris (pH 7.4), 200 mM NaCl, 200 mM KCl, 2 mM CaCl_2_, 0.01% sodium azide and 0.1% low-viscosity sodium alginate (Sigma-Aldrich, A1112)) in a 384-well UV-STAR® microplate (Greiner Bio-One International). Alginate lyase activity was determined by measuring the increase in absorbance at 235 nm at one minute intervals using a Tecan Spark Multimode Microplate Reader (Tecan Inc.).

---

**Figure S1:**
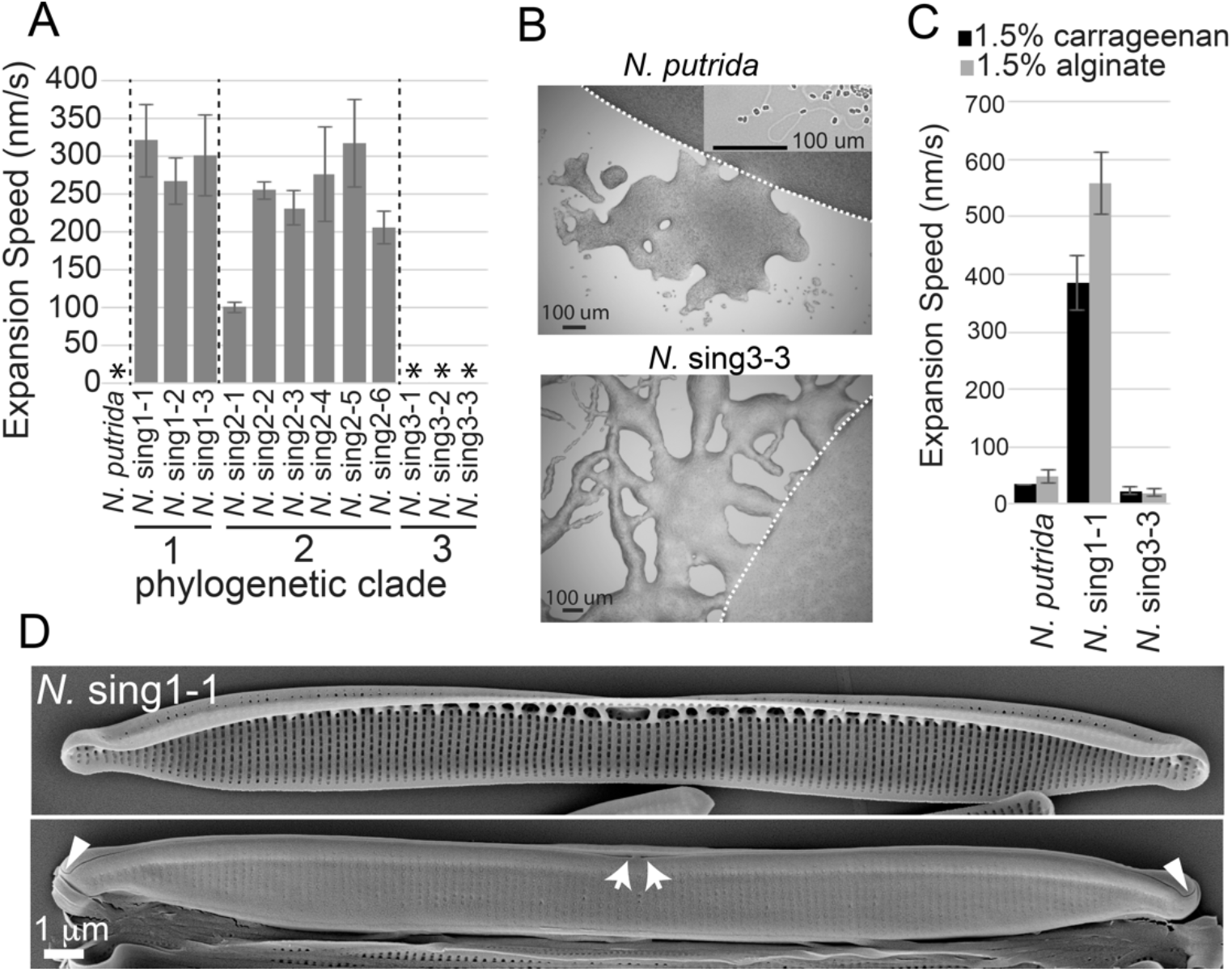
Growth and scanning electron microscopy of apochlorotic diatoms. **(A)** The rate of radial colony expansion on seawater agar medium is shown for the indicated diatom isolates. Asterisks mark isolates growing too slowly to be measured by this method. **(B)** The images show the colony periphery of the indicated diatoms grown on seawater agar medium. The dashed white line marks the perimeter of the primary colony. Note that while gliding motility is minimal, a degree of motility is indicated by multicellular networks of cells that have migrated away from the primary colony. Scale bar = 100 μm. Note that *N. putrida* occasionally makes sectors containing dispersed cells and EPS trials that can be seen by bright field microscopy (inset). **(C)** Quantification of motility for the indicated species. **(D)** Scanning electron microscopy (SEM) of *N.* sing1-1 valves. The upper and lower panels show internal and external views, respectively. White arrows point to the proximal raphe. Arrowheads point to hooks at the distal raphe ends. Scale bar = 1 μm.

**Figure S2:**
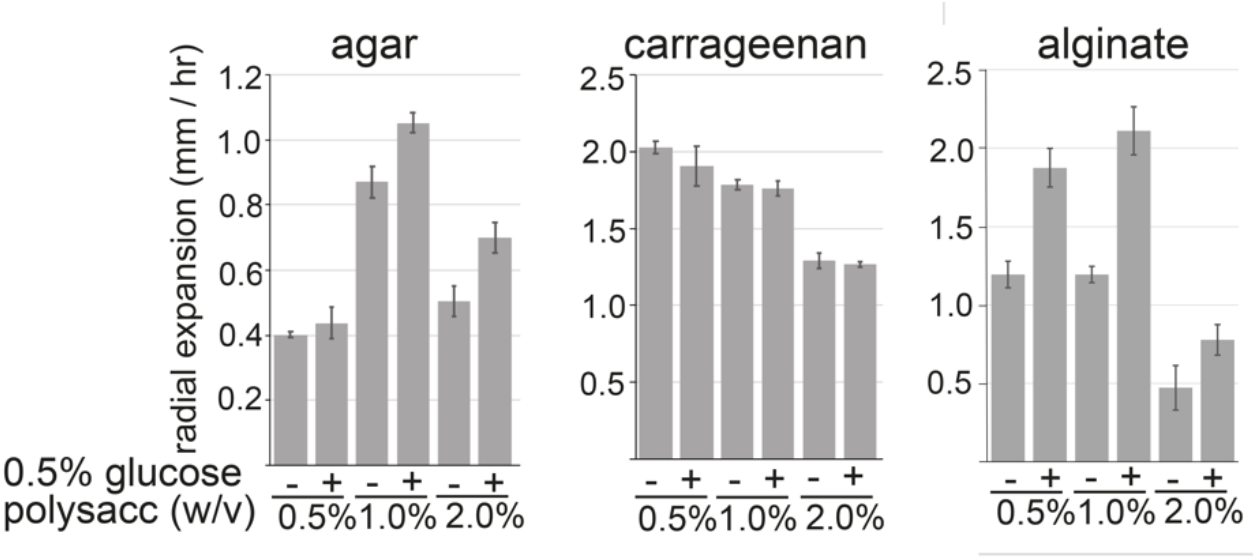
Measurement of rates of radial colony expansion. The graphs show the speed of *N*. sing1-1 radial colony expansion on the indicated polysaccharides in the presence (+) or absence (-) of 0.5% glucose. The measurements were made in triplicate. Standard deviation is indicated.

**Figure S3:**
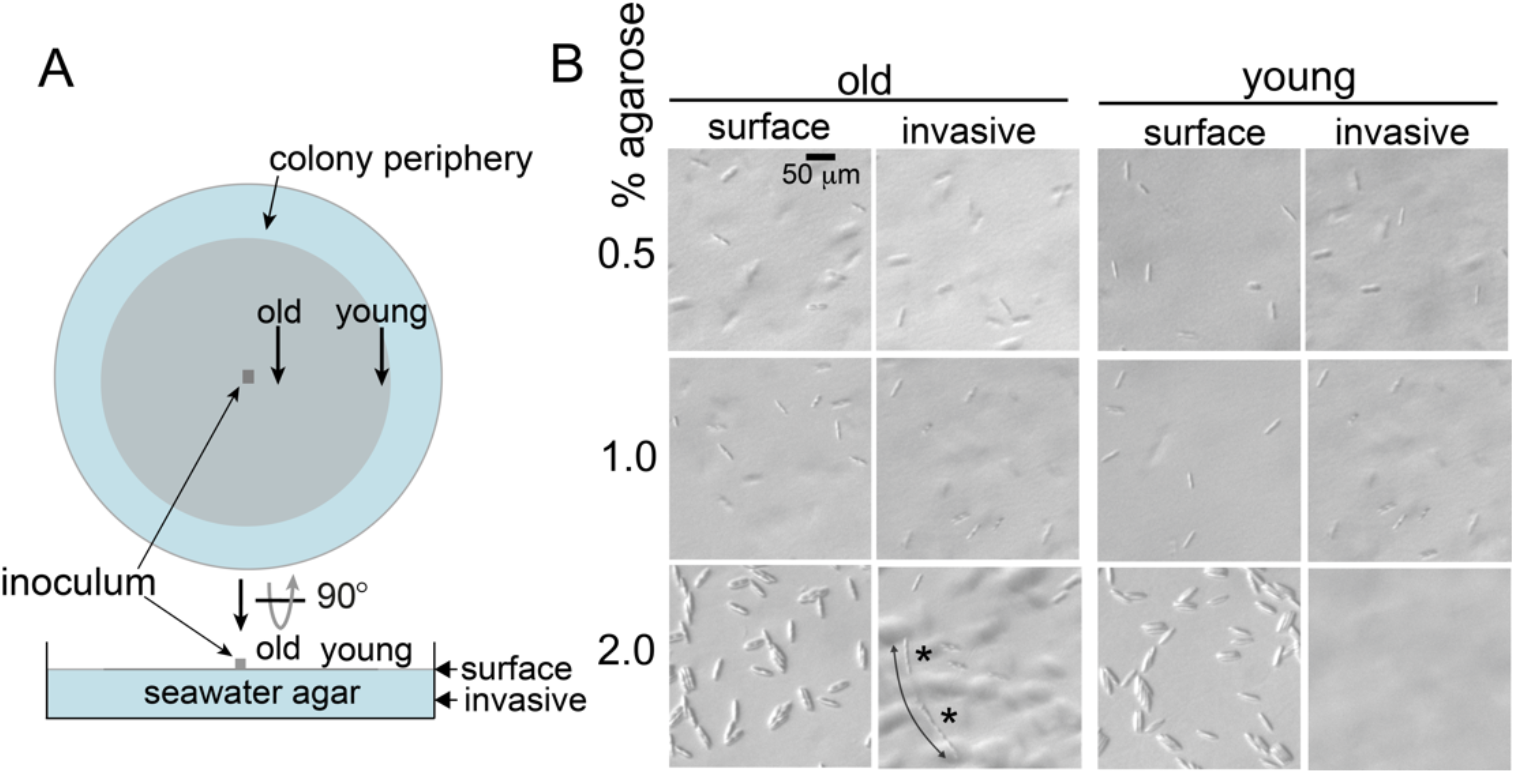
Invasive growth of *N.* sing1-1. **(A)** Diagram showing how invasive growth is documented. The seawater agar is inoculated at the center of the petri dish and then imaged at young and old regions as indicated. Diatoms are imaged on the surface of the medium (surface) and at a depth of approximately 2 to 3 mm (invasive). **(B)** Diatoms growing on the surface and invasively on the indicated media. Note that at high agar concentrations (2%), diatoms are seen growing invasively in old but not younger regions of the colony. Those growing invasively can be seen in chains of cells (asterisks) that appear to be moving in the same channel (double-headed arrow). Scale bar = 50 μm.

**Figure S4:**
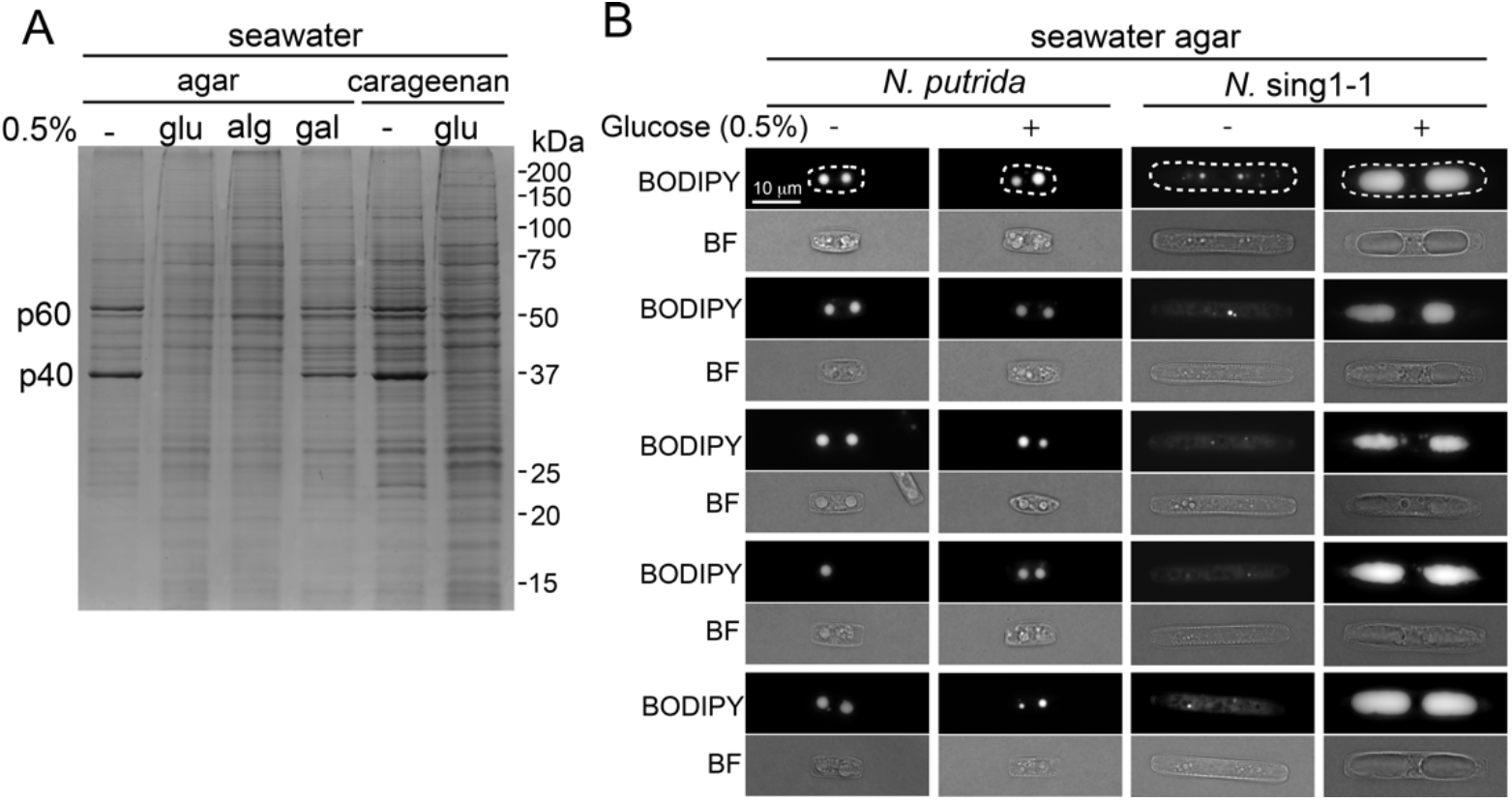
Catabolite repression and lipid droplets. **(A)** SDS-PAGE of total cell extracts from *N.* sing1 diatoms grown on seawater agar or seawater carrageenan medium in the presence (+) or absence (-) of 0.5 % (w/v) of the indicated carbohydrates. glu, glucose; alg, alginate; gal, galactose. Catabolite-repressed proteins p40 and p60 are identified. Related to Figure 2C. **(B)**. The diatoms grown on seawater agar medium in the presence or absence of 0.5% glucose. Lipid droplets were stained with BODIPY as described in the materials and methods. Cells are shown by fluorescence (BODIPY) and bright-field (BF) microscopy. Five randomly selected cells are shown for each condition. The dashed white boxes outline the cells shown in Figure 2D. Scale bar = 10 μm. Related to figure 2D.

**Figure S5:**
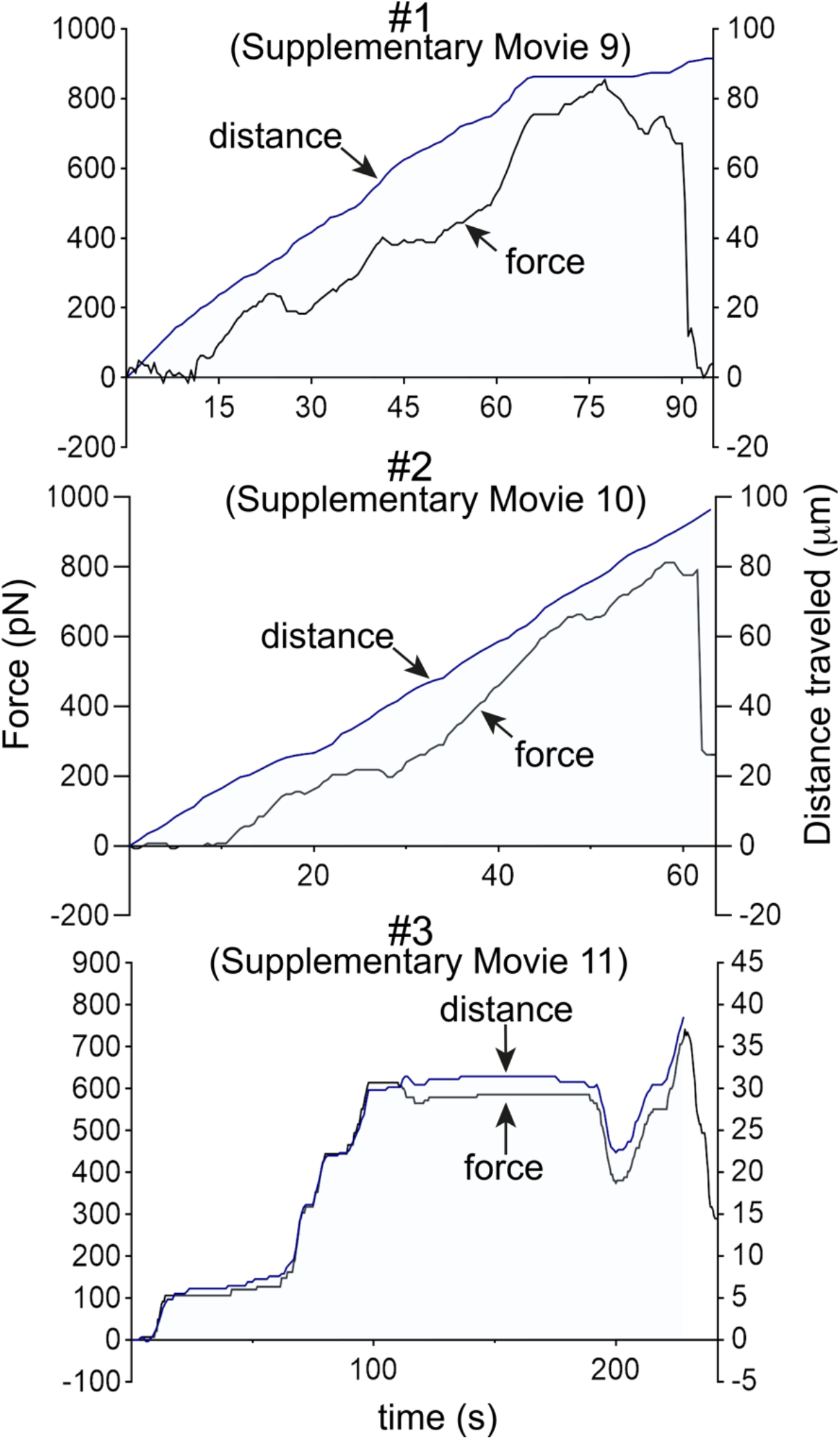
Force/velocity graphs from micropipette experiments. Force from microneedle deflection is plotted along with distance traveled by the diatom. Distance (blue line) and force (black line) are shown. Data are from diatom measurements #1, #2 and #3 shown in Figure 6B. Related to Supplementary Movies 9, 10 and 11.

**Figure S6:**
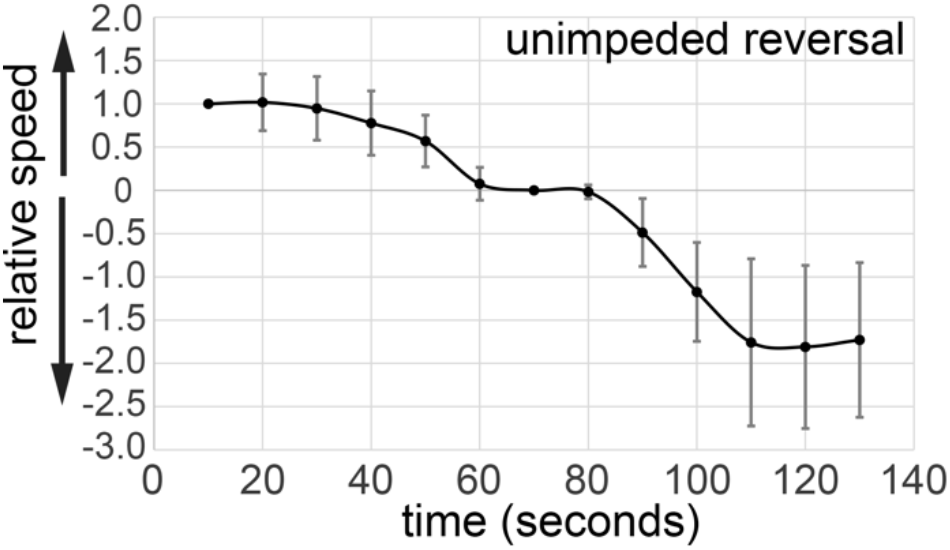
Measurement of free-running reversal. The graph shows the relative speed of unimpeded diatoms undergoing reversal. Note that they gradually slow before stopping. The black dot shows the mean (n=20). Standard deviation is indicated. Related to Figure 6C.

**Figure S7:**
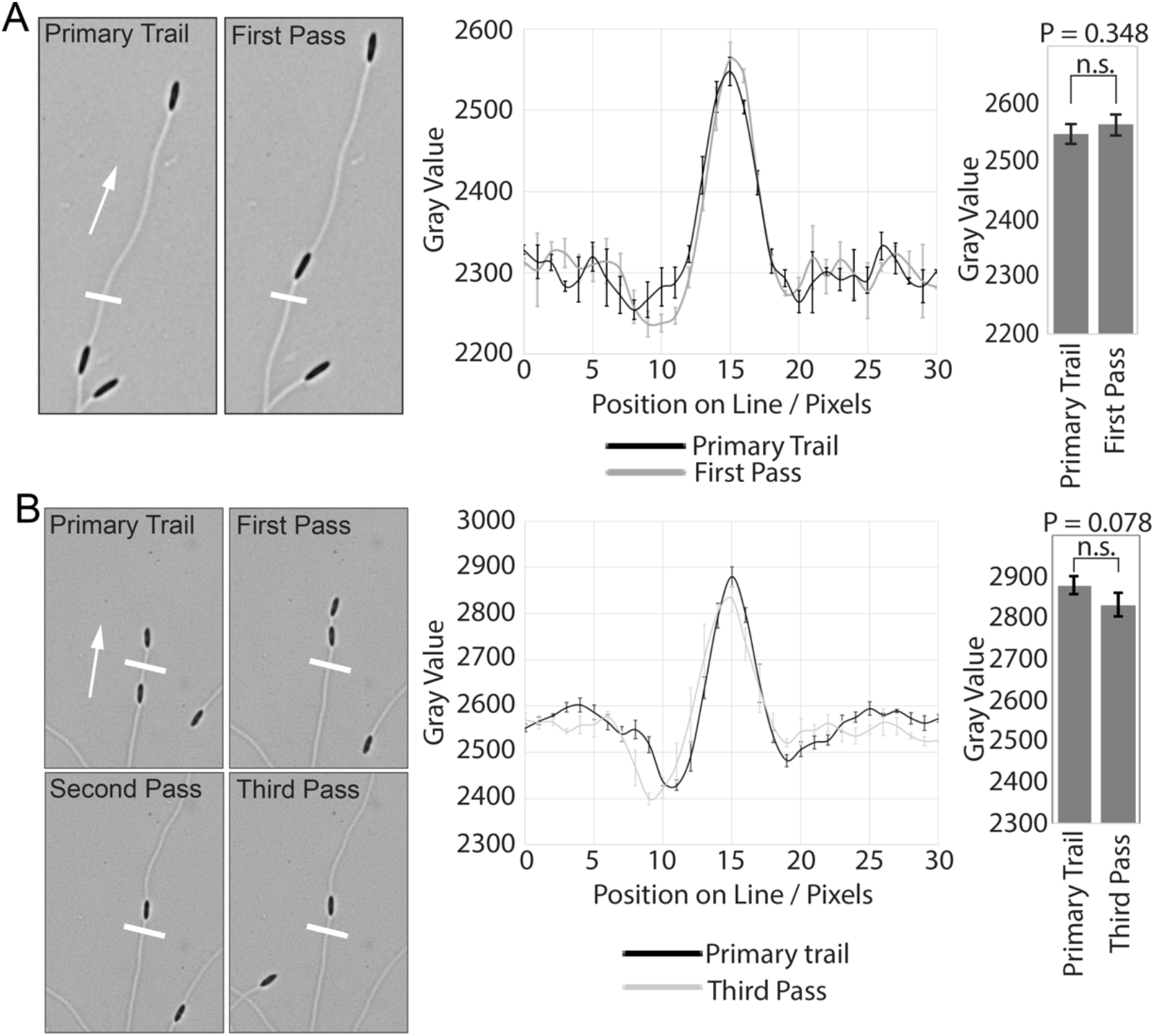
The appearance of EPS trails made by blazers does not change dramatically with the passage of runners. **(A)** Documentation of a primary trail that was passed over once. **(B)** Documentation of a primary trail that was passed over three times. The left panels show images of trails. The panels are labeled according to how many times the primary trail had been passed over by additional diatoms. The white line indicates the position of line scans, which are shown quantified for gray value in the middle graph. The graph on the right compares the peak gray values at the indicated points in time. Note that there is no significant difference in peak gray values.

**Figure S8:**
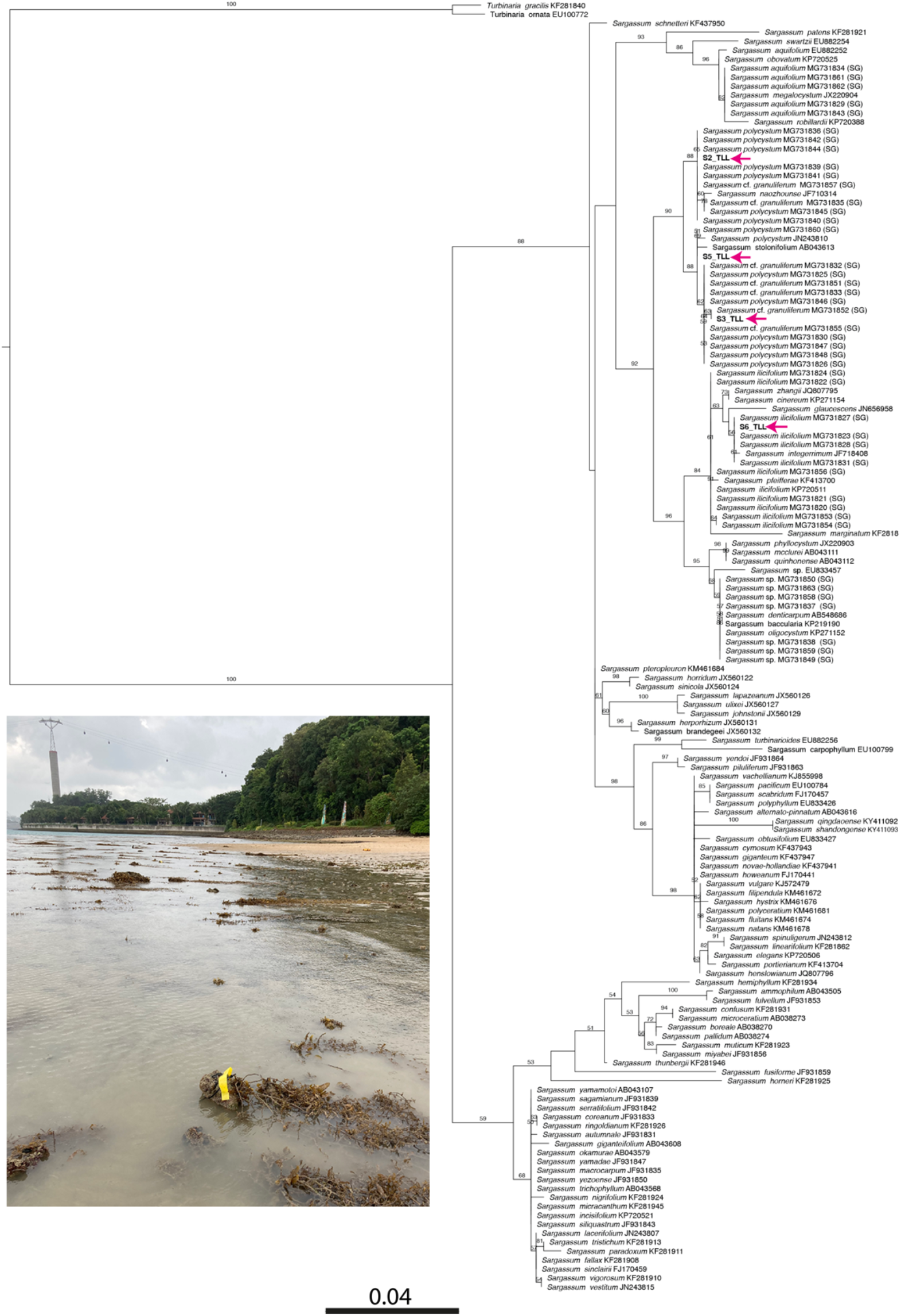
Identification of *Sargassum* species. Phylogenetic reconstruction of *Sargassum* (Phaeophyceae) with *Turbinaria* as outgroup, based on a maximum likelihood (ML) analysis of ITS-2 alignment. Bootstrap proportions are indicated for maximum likelihood when >0.5. Four *Sargassum* isolates from which diatoms were recovered are shown in bold and identified with a magenta arrow. The inset photograph shows *Sargassum* (yellow tag) at the Sentosa, Singapore collection site.

## Notes

### Competing Interest Statement

The authors have declared no competing interest.

### Summary of Updates

minor edits

